## Supplementary figure for "Signal-based optical map alignment"

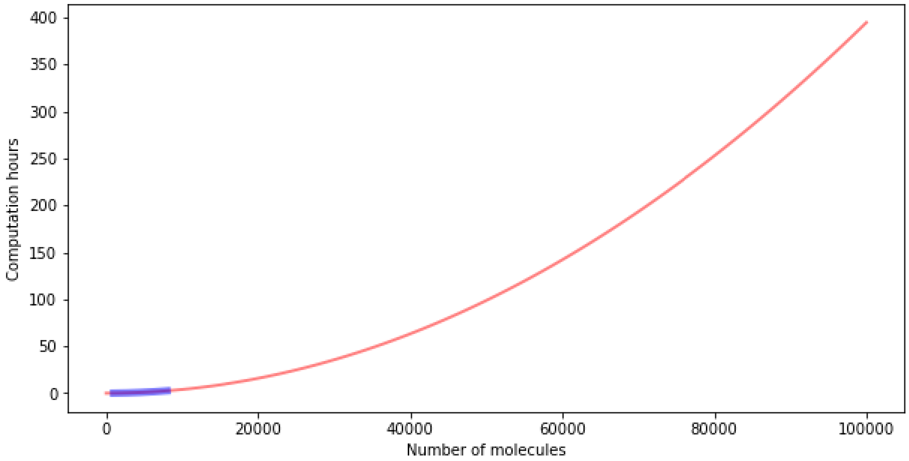


Figure S1: **Estimated time required to compute of all-vs-all OptiMap-naive alignments.** Obtained by fitting a quadratic function $f(x) = mx^2$ on run-times calculated for datasets consisting of 0 to 8,000 molecules with a step of 200. The optimal value of $m$, obtained using non-linear least squares, was $1.34 \times 10^{-4}$ with a standard deviation of $8.8 \times 10^{-7}$.
